# Evaluating methods for B-cell clonal family assignment

**DOI:** 10.1101/2024.05.29.596491

**Authors:** Katalin Voss, Katrina M. Kaur, Rituparna Banerjee, Felix Breden, Matt Pennell

## Abstract

The adaptive immune response relies on a diverse repertoire of B-cell receptors, each of which is characterized by a distinct sequence resulting from VDJ-recombination. Upon binding to an antigen, B-cells undergo clonal expansion and in a process unique to B-cells the overall binding affinity of the repertoire is further enhanced by somatic hypermutations in the receptor sequence. For B-cell repertoires it is therefore particularly important to analyze the dynamics of clonal expansion and patterns of somatic hypermutations and thus it is necessary to group the sequences into distinct clones to determine the number and identity of expanding clonal families responding to an antigen. Multiple methods are currently used to identify clones from sequences, employing distinct approaches to the problem. Until now there has not been an extensive comparison of how well these methods perform under the same conditions. Furthermore, since this is fundamentally a phylogenetics problem, we speculated that the mPTP method, which delimits species based on an analysis of changes in the underlying process of diversification, might perform as well as or better than existing methods. Here we conducted extensive simulations of B-cell repertoires under a diverse set of conditions and studied errors in clonal assignment and in downstream ancestral state reconstruction. We demonstrated that SCOPer-H consistently yielded superior results across parameters. However, this approach relies on a good reference assembly for the germline immunoglobulin genes which is lacking for many species. Using mPTP had lower error rates than tailor-made immunogenetic methods and should therefore be considered by researchers studying antibody evolution in non-model organisms without a reference genome.

## Introduction

B-cells and their diverse repertoires of receptors are a central component of the adaptive immune response. Naive B-cells, which have not previously encountered foreign antigens, can become activated upon binding of their B-cell receptors (BCRs) to antigens presented by pathogens. Upon activation, these B-cells undergo proliferation and differentiation, ultimately leading to the secretion of antibodies specifically designed to recognize and bind the encountered pathogens. These antibodies play a crucial role in the immune defense by either directly neutralizing pathogens or triggering downstream immune responses that lead to pathogen clearance. A diverse repertoire of BCRs is necessary to recognize a broad spectrum of pathogens. This diversity is achieved through two primary mechanisms (Figure 1A): VDJ-recombination and somatic hypermutations (SHM). B-cell receptors are composed of two identical heavy chains and two identical light chains. For this study, we concentrate on the heavy chain. The heavy chain locus encompasses V, D, and J genes, and through VDJ-recombination, one V gene, one D gene, and one J gene are joined together. In humans we know of 44 functional V genes, 27 functional D genes and 6 function J genes [1]. Consequently the VDJ-recombination contributes significantly to the vast diversity observed in B-cells. Another big contributor to the diversity of the BCRs is the addition or removal of P and N nucleotides at the junctions of the genes during VDJ-recombination [2]. The parts of the BCRs that bind to antigens are called complementarity-determining regions (CDRs). There are 3 CDRS: CDR1 and CDR2 are encoded in the V-gene, the CDR3 region encompasses part of the V-gene, the P and N nucleotides, the D gene and part of the J gene, and is a strong determinant of the specificity of each receptor. Following antigen binding, a B-cell undergoes affinity maturation, a process characterized by clonal expansion and SHM: The B-cell proliferates into clones and SHM introduce point mutations to the B-cell receptor sequences. These mutations enhance antibody diversity and can lead to the production of antibodies with increased affinity for the antigen [2]. A clonal family refers to the collective group of B cells originating from a single VDJ rearrangement event.

**Figure 1:**
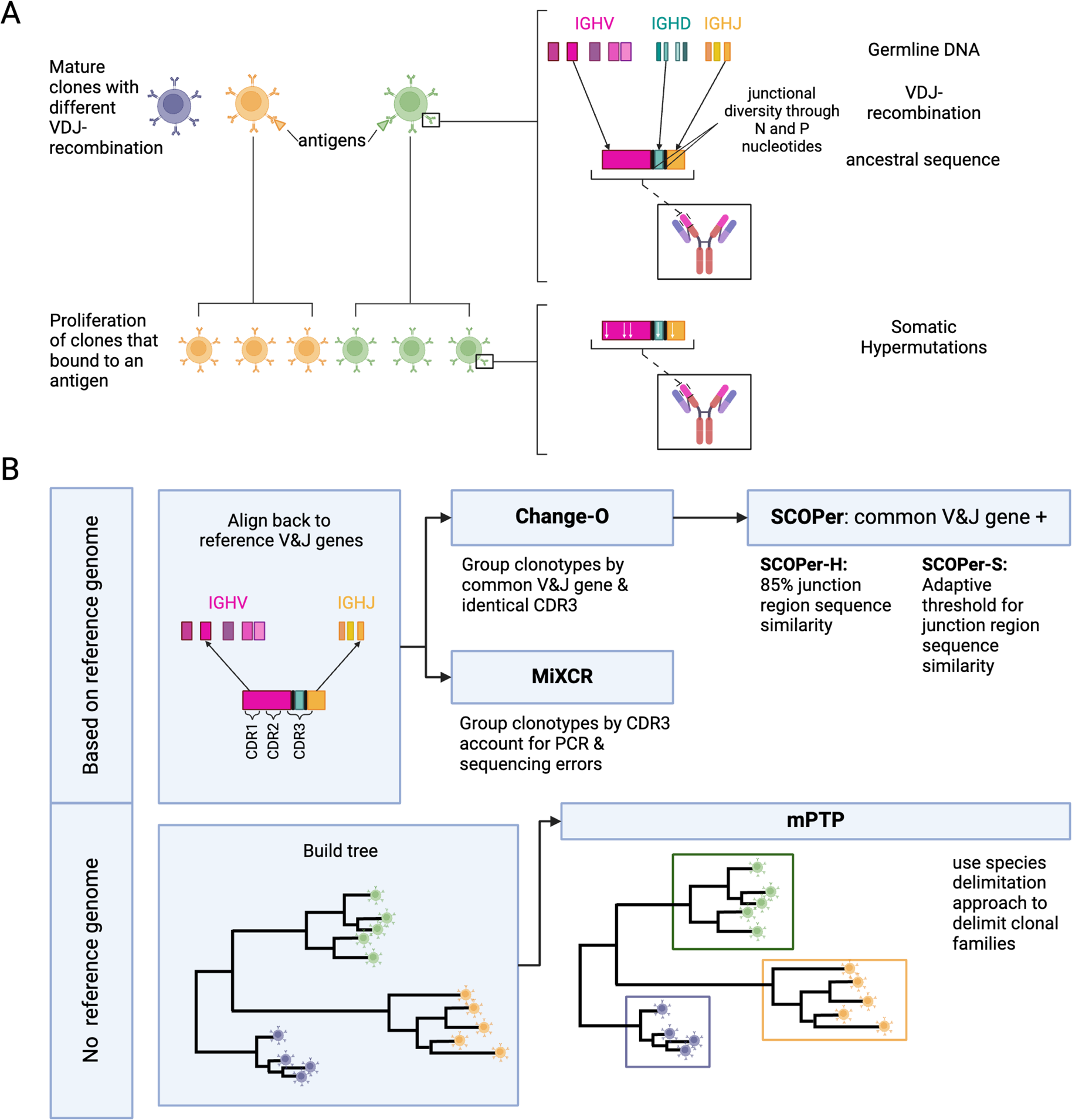
A) Development of B-cell receptors and their sequences. Mature B-cell clones stem from B-cell precursors. Their BCR sequences arise through VDJ-recombination on the light and heavy chain. During VDJ-recombination N and P nucleotides are added in between the junctions leading to more junctional diversity. Once a mature B-cell binds to an antigen it proliferates into more clones. During proliferation the rearranged IG genes undergo somatic hypermutations. **B) Overview of the methods and their requirements**. We used two types of methods: methods dependent on a reference genome and methods independent of a reference genome. For the independent method, mPTP [15], we first build a phylogenetic tree of all sequences with RAxML-NG [16]. mPTP then fits two Poisson Processes, one for speciation and one for coalescence, to the data and groups the clones accordingly. The reference-genome-dependent methods first align the sequences to a reference genome to find the used V and J genes. This alignment is either done by the methods themselves (MiXCR [8]) or done through IMGT [17]. MiXCR delimits the clones by the CDR3 region, accounting for PCR and sequencing errors. Change-O [9] delimits the clones by a common V and J gene and an identical CDR3 region. SCOPer [10] uses the common V and J groups provided by Change-O. The hierarchical method then further groups by 85% junction region similarity. The spectral method calculates an adaptive threshold for junction region similarity.

The human body harbors approximately 10^11^ B-cells [3], suggesting a vast array of clonal families. Recent advances in high-throughput sequencing technology have revolutionized the field of BCR repertoire sequencing [4], enabling the analysis of the clonal relationships of BCRs. One of the central challenges in B-cell analysis lies in accurately delineating these clonal families within sequencing data from each individual. As the sequences within a family originate from the same ancestral B-cell, they should not be treated independently in statistical analysis. Only once the clonal families have been identified, is it possible to infer which ones have expanded in response to antigen binding. Subsequent analyses can include examination of VDJ gene usage, calculation of somatic hypermutation (SHM) statistics, quantification of selection during affinity maturation [5], and inference of the original receptor sequence and identification of the original antigen target. Furthermore, by tracing the development and diversification of B-cell lineages, it is feasible to identify the specific genetic and structural alterations that give rise to antibodies capable of neutralizing a wide spectrum of pathogenic strains. This identification of broadly neutralizing antibodies serves as the foundation of effective vaccine design against challenging pathogens such as HIV [6, 7].

Multiple strategies to delimit B-cell clones are currently employed. In this investigation we will focus on four state-of-the-art methods that all rely on a reference genome, MiXCR [8], Change-O [9], SCOPer hierarchical (SCOPer-H) [10], and SCOPer spectral (SCOPer-S) [11], and one that analyzes the unannotated sequences in a phylogenetic context, mPTP. MiXCR involves an initial alignment of sequences to a reference genome, followed by the assembly of clonotypes based on identical sequences for user-defined gene features like the CDR3 region since it encompasses the majority of the diversity of BCR sequences. By allowing fuzzy matches MiXCR tolerates PCR and sequencing errors [8]. Change-O requires a preceding alignment performed by IMGT/HighV-QUEST [12], IgBLAST [13] or iHMMune-align [14]. Subsequently, Change-O utilizes these alignments to reconstruct germline sequences and proceeds to group the sequences by common V gene, J gene, and junction region. The junction region is defined as the CDR3 region plus flanking amino acid residues [11]. SCOPer, an extension of the Change-O toolkit, was developed by the same research group, and it is specifically designed for the assignment of B cell clones. In our study, we employed two models of SCOPer: the hierarchical model (SCOPer-H) and the spectral model (SCOPer-S). Both of these models utilize the outcomes generated by Change-O as their input. The approach of SCOPer-H uses the assumption that sequences sharing highly similar junction regions likely originate from the same clonal ancestor, since it is unlikely that different recombinations result in the same junction region. Therefore it uses a user-defined cutoff delineating the minimum similarity threshold for two junction regions to be considered clonally related. In contrast to SCOPer-H, SCOPer-S takes an adaptive approach by calculating the optimal cutoff for each instance [11]. A comparison of the tools’ requirements can be found in Table 1.

**Table 1:** Comparison of the different tools. All presented methods have distinct preprocessing procedures and employ varied methodologies to address the clonal family assignment problem. Here we list the specific requirements of each tool and the different approaches to this problem.

|  | MiXCR | Change-O | SCOPer-H | SCOPer-S | mPTP |
| --- | --- | --- | --- | --- | --- |
| aligns sequence to reference genome | ✓ | ✓ | ✓ | ✓ |  |
| creates groups based on common V and J gene |  | ✓ | ✓ | ✓ |  |
| compares CDR3/junction region between sequences | ✓ | ✓ | ✓ | ✓ |  |
| uses a threshold for sequence similarity | ✓ |  | ✓ | ✓ |  |
| requires a tree |  |  |  |  | ✓ |

The clonal family assignment is similar to the species delimitation problem in phylogenetics: Phylogenetic trees representing multiple species exhibit similarities to trees representing B-cell clones. Instead of delimiting species that have evolved over thousands of years, the objective shifts to delimiting clones that have developed over shorter time frames, typically days or weeks. We realized that phylogenetic tools originally designed for species delimitation could offer an alternative approach to address the clonal family assignment problem. This approach offers distinct advantages, particularly in scenarios where reliance on a reference genome is impractical or challenging, such as when working with species lacking a reliable reference genome. For non-model organisms reference genomes are based on a much smaller sample size, which can lead to missing or false alleles. Even though recent advances in sequencing have helped to sequence many germline Ig repertoires of organisms that have not been sequenced before [18], the annotation of B-cell receptors for their V, D and J genes remains a challenge [19]. Consequently, methods reliant on a reference genome may encounter greater difficulty in accurately discerning clones compared with those that operate independently of a reference genome.

One such approach is the multi-rate Poisson Tree Processes (mPTP) method [15]: This method delimits single-locus species, or in our case clones, given a binary phylogenetic tree. mPTP builds upon the foundation of the simpler Poisson tree processes (PTP) approach [20]. PTP assumes a model comprising two Poisson processes: one representing speciation events and the other modeling the coalescent process. mPTP further refines the PTP model by introducing a nuanced adaptation: instead of fitting a single branching rate for all coalescent events, mPTP assigns a distinct branching rate for each species. Applying this assumption to our clonal family problem, one Poisson process should represent VDJ-recombination (analogous to speciation) and another Poisson process should represent SHM (diversification within a clone). Furthermore, each clone is expected to exhibit varying SHM rates, reflecting the diverse evolutionary trajectories of individual clonal families. These assumptions align with those of SCOPer-S, which also accounts for SHM variation among different clonal families. Therefore, we are particularly interested in comparing the performance of these two tools that assume variable evolutionary rates across clonal families, SCOPer-S and mPTP. If mPTP demonstrates a competitive performance, it would present a valuable alternative for analyzing sequences from organisms lacking a reliable reference genome (Table 1). In other contexts mPTP has demonstrated successful application across various experimental datasets. In the study conducted by Kapli et al., mPTP was applied to analyze 24 different genera using the COI-5P gene, resulting in superior performance compared to other tools [15]. Furthermore, mPTP has been employed in independent studies on free-living amoebae [21], Plecoptera [22], radicine pond snails [23], and freshwater mussels [24]. These studies collectively highlight the versatility and effectiveness of mPTP across diverse biological datasets.

We raise the unanswered question: What strategy is the best to discern clones in B-cells among current state-of-the-art methods and a native phylogenetic strategy? By conducting simulations of B-cell repertoires focused on the heavy chain, considering variables such as clone count, SHM, and average lineage count per clone, we aim to comprehensively measure and compare the performance of the state-of-the-art tools in B-cell analysis as well as a phylogenetic method. An overview of the methods and a visualization of our pipeline is shown in Figure 1B. We adopted a multifaceted approach to assess the performance of each method, employing measures such as the Mean Squared Error (MSE) of the median family size, the number of discerned clonal families, and the F1-score. The MSE and the number of identified families offer insights into overall trends, while the F1-score, being the harmonic mean of precision and recall, provides a detailed understanding of method performance and serves as our primary performance metric. Additionally, we investigated the impact of the tools’ performance on downstream analysis, particularly focusing on ancestral sequence reconstruction. This serves as a standardized foundation for future studies delving into B-cell data analysis, providing valuable insights into optimal tool selection under various conditions.

## Results and Discussion

### SCOPer-H outperforms all other methods across variety of conditions

In our evaluation we considered multiple measures across the parameters and tools. For all tools except mPTP, an increase in the SHM rate resulted in fewer sequences being analyzed (Figure S1). We classified the missing samples as singletons— clonal families with only one sequence. To mitigate the distortion caused by singletons, which are often disregarded in real-life analysis [25], we excluded all singletons from our analysis. Initially, we assessed the MSE of the median family size for all tools. On average, SCOPer-H exhibited the best performance based on the MSE metric (Figure S2). To further interpret this finding, we analyzed the number of clonal families identified by each method compared with the actual number of families. Across all methods we observed a consistent pattern of overestimating the numbers of clonal families, even after removing the discerned singletons as seen in Figure 2. This indicates a tendency for all methods to oversplit clonal families. SCOPer-H gets closest to the correct number of clonal families across all leaf and SHM configurations. SCOPer-S has the second best performance on this measure in most leaf configurations, but is outperformed by mPTP for the three lowest SHM values. It is notable that on average SCOPer-S has the smallest interquartile range whereas mPTP has the largest. MiXCR and Change-O perform similarly to each other and poorly relative to the other tools.

**Figure 2:**
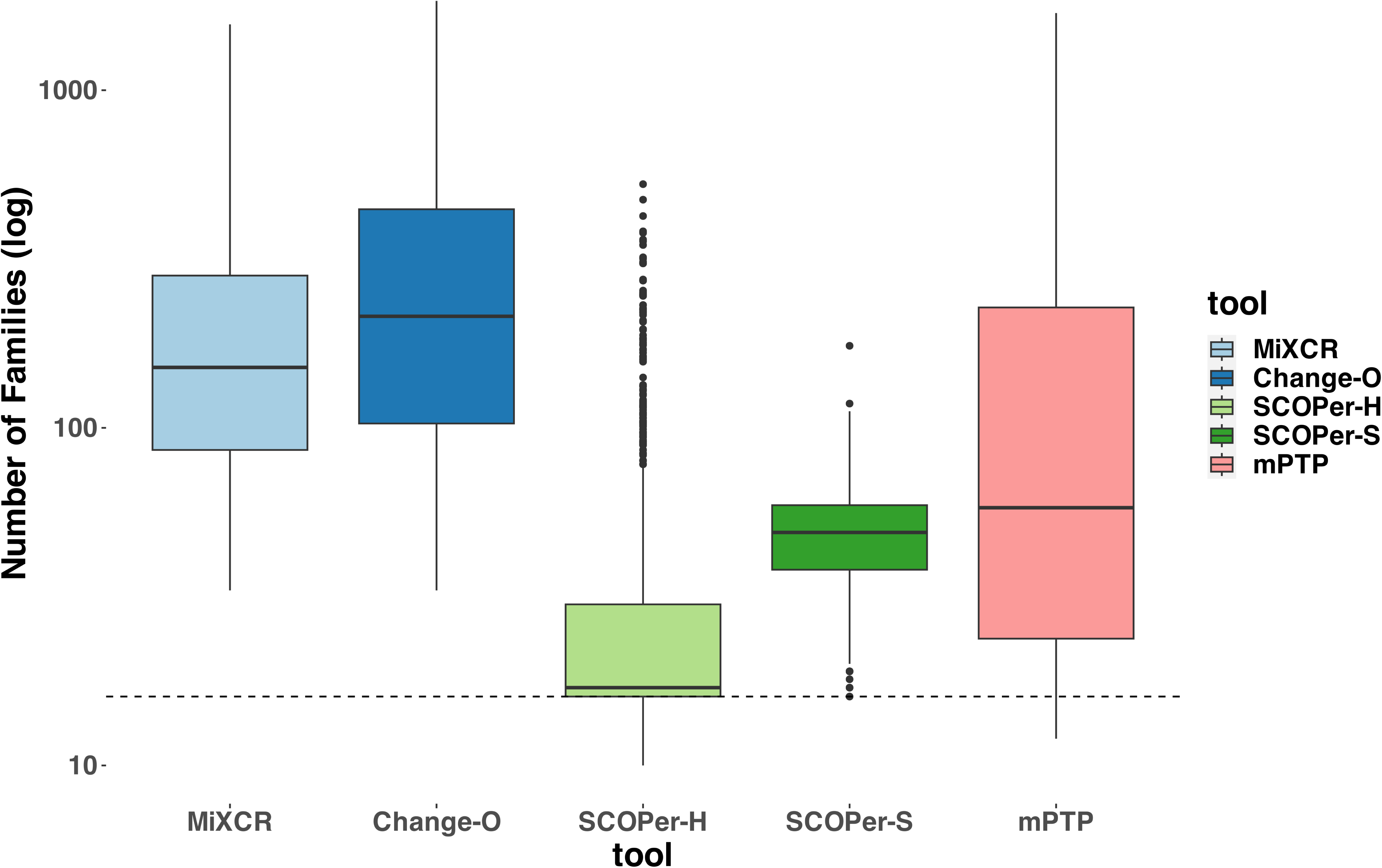
Number of Clonal Families (log scale) discerned by the different tools. For this analysis we removed discerned singletons. We counted the number of clonal families that the methods discerned for each simulation and compared it to the correct number of clonal families (in this case 16). The dashed line represents the correct number of clonal families. We can clearly see that all methods consistently return a higher number of families, indicating the subdivision of one correct family into multiple. SCOPer-H on average performs the best.

To provide a more nuanced assessment of the tools’ performances, we calculated the F1-score, which represents the harmonic mean of precision and recall. Across all parameter configurations, SCOPer-H consistently outperformed all other methods by a substantial margin (Figure 3, Figure S3). mPTP and the spectral model emerged as contenders for the second-best performance. SCOPer-S exhibited superior performance at higher leaf and SHM configurations, whereas mPTP demonstrated better performance at lower leaf and SHM configurations (Figure S3). MiXCR and Change-O again demonstrated the poorest performances, with MiXCR slightly outperforming Change-O. We have similar explanations for the poor performance of both MiXCR and Change-O: MiXCR groups sequences solely based on identical matches and then allows for fuzzy matches to accommodate PCR and sequencing errors. However, the SHM rate is high and likely surpasses what MiXCR’s fuzzy matching can accommodate. As a result, MiXCR tends to oversplit clonal families. Similarly, Change-O initially groups sequences by VJ-genes and then by identical junction regions, failing to group sequences from the same clonal family with SHM in the junction region. Consequently, Change-O also tends to oversplit clonal families. The same patterns could be observed across all numbers of clones that we simulated in a separate simulation set (Figure S4).

**Figure 3:**
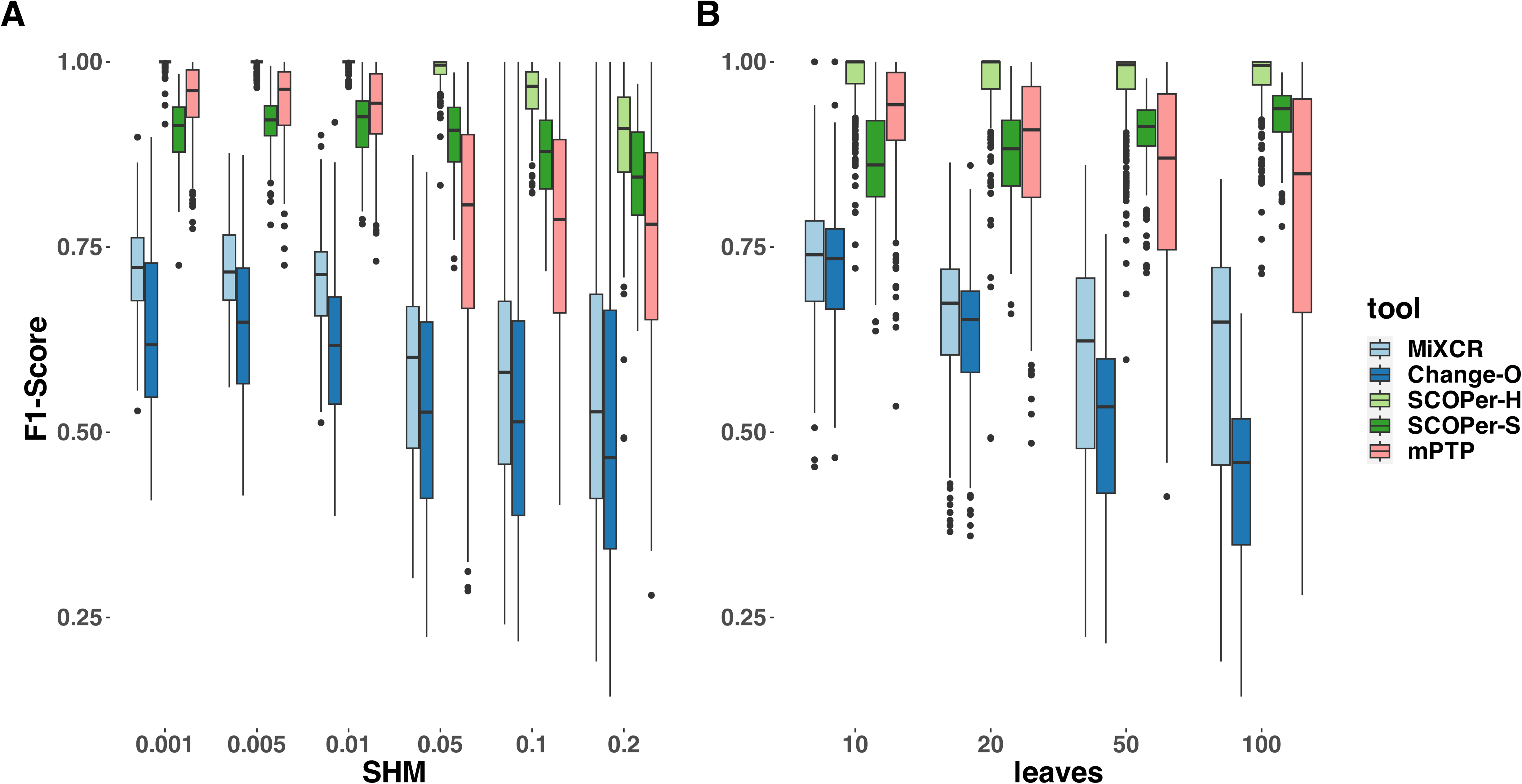
F1-Score yielded by the different methods. For this analysis we removed singletons. A) different SHM rates B) different average number of leaves per clonal family. The F1-score is the harmonic mean of precision and recall, a score of 1 meaning perfect precision and recall.

**Figure 4:**
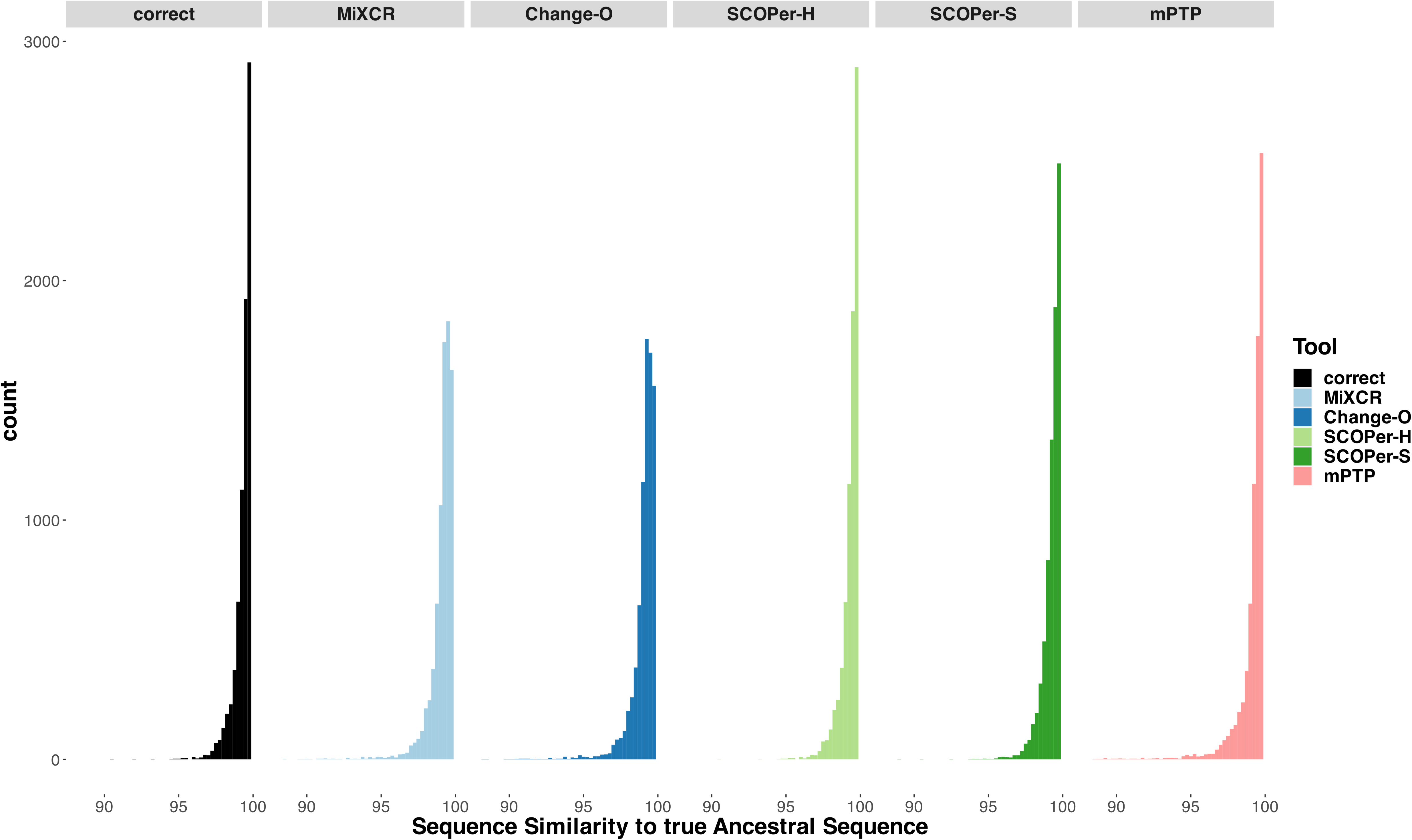
Sequence Similarity between the real ancestral sequence and the derived ancestral sequence based on the clonal families discerned by the methods. If all 5 methods discerned a specific family the only difference being the amount of sequences in a family, we considered it in this analysis. The “correct” column acts as a point of comparison, using the correct sequences of the clonal family for ancestral sequence reconstruction.

Our analysis clearly identified SCOPer-H to be the best method in our selection of tools across parameters. We explored various thresholds for sequence similarity which revealed a consistent trend: higher thresholds led to improved performance for SCOPer-H (Figure S5). Across the range of thresholds tested, all options yielded superior or comparable results compared to SCOPer-S. This contradicted the anticipated superiority of SCOPer-S over SCOPer-H. SCOPer-S was designed to enhance accuracy by dynamically calculating an optimal threshold for similarity within each VJ-group obtained from Change-O, rather than employing a fixed threshold for all groups. However, our findings deviate from the anticipated outcomes and contradict the results of the tool’s authors [10]. To delve deeper into this discrepancy, we designed a simulation set specifically targeting the junction region, as this is the focal point of SCOPer’s analysis for both models. We incorporated the parameter “junction region length” into our setup, considering the developers’ indication that the performance of the hierarchical model depends on the junction region length, which we had not analyzed in our previous simulations. For these simulations we randomly sampled V,D and J genes from the ImMunoGeneTics (IMGT) [17] reference directory and joined them together, adding a varying amount of nucleotides in between. We also only simulated SHM at the junction region, to analyze the effect it has on the performance of SCOPer-H. Analysis of the F1-score revealed a notable decline in performance for SCOPer-H at an SHM rate of 0.2. Further exploration pinpointed this decline to junction region lengths of 70 and above (Figure S6). We were not able to replicate the finding that the performance of SCOPer-H declines for shorter junction regions [10], and in most of our new simulations SCOPer-H still outperformed SCOPer-S (Figure S6).

To validate our findings and ensure they were not biased by our simulations, we also ran both models on subsets of the simulation data provided by Nouri et al. [11]. The results confirmed that SCOPer-H generally outperforms the spectral model in typical scenarios (Figure S7). In Nouri et al.’s comparison of the spectral and hierarchical model their validation primarily relied on a limited simulation setup and real-life data [10]. In their validation process using real data, the emphasis was on confirming highly homogeneous discerned clonal families. This favors SCOPer-S because of its tendency to oversplit clonal families, resulting in each discerned family being highly homogeneous. Although they later conducted an extensive simulation, they only evaluated the performance of SCOPer-S. Our simulations revealed that SCOPer-S is overly stringent, resulting in the oversplitting of clonal families (Figure 2,3,S7).

### mPTP as a new alternative not reliant on a reference genome

Across a majority of parameters we demonstrated that mPTP outperforms all other methods but SCOPer-H. Particularly noteworthy is its superior performance compared with SCOPer-S, which shares a similar approach to mPTP but relies on additional information from a reference genome. For low SHM rates, mPTP has the second best performance on average across all tools. This is particularly striking because it is the only method that does not require any information except the sequences. To explore the SHM rates in a more biologically meaningful context, we calculated the distance of the sequences to their ancestral sequences. In nature, there can often be a divergence of up to 5% between antibody sequences and their original germline sequence. In our simulations, sequences diverge on average less than 5% from their germline sequence for SHM rates of 0.001, 0.005, 0.01, and 0.05 (Figure S8). This indicates that mPTP performs well in scenarios that are biologically realistic. As mPTP appears to be a promising alternative in our simulation setting, we aimed to evaluate its performance in situations where methods dependent on a reference genome fail. To do so, we assessed the performance of the tools on a dataset where a reliable reference genome is not available. We achieved this by creating a simulation set with “fake” V genes, by introducing insertions, deletions, and mutations to the known V gene sequences from IMGT. We then applied all methods to this simulation set using the same parameters as before. We observed that all the tools relying on a reference genome did not return a substantial number of input sequences in their results (Figure S9). While we had already noticed a pattern of missing samples with an increase in the SHM rate in the original simulations (Figure S1), this pattern was magnified with the fake V genes. mPTP, not being dependent on a reference genome, consistently returned all input sequences (Figure S9). To evaluate the overall performance, we classified the missing samples as singletons and calculated the F1-Score with singletons included. For the three lower SHM rates, mPTP outperforms all other methods (Figure S10). However, starting at a SHM rate of 0.05, the performance of mPTP decreases significantly. This pattern was also observed in the original simulations (Figure 3). By examining the number of singletons per simulation, we found that this decrease in performance is due to over splitting (Figure S11). At a SHM rate of 0.05 or higher, mPTP struggles to correctly differentiate between sequence differences caused by different VDJ recombination events and those caused by SHM. Our analysis revealed mPTP to be a valuable alternative to other methods, particularly when the organism of interest lacks a robust reference genome. This makes mPTP a valuable tool for analyzing B-cell repertoire data in diverse contexts, including species with poorly characterized genomes or non-model organisms. Consequently, researchers can leverage mPTP to gain insights into clonal relationships and dynamics without being hindered by limitations associated with reference genome availability or quality.

### Ancestral Sequence Reconstruction

The reconstruction of the ancestral sequence from sequences within clonal families is a vital aspect of repertoire analysis. It helps to determine the naive sequence responsible for initially binding to a specific antigen, and in some cases what that initial stimulating antigen even was. We wanted to evaluate the extent to which errors in clonal family assignment impact ancestral sequence reconstruction, a common downstream inference. For each method, we assessed the ability to reconstruct ancestral sequences from all identified clones comprising more than two sequences. This reconstruction process relied on phylogenetic trees constructed from the sequences of each discerned family. We used RAxML-NG [16] for constructing all trees and for reconstructing the ancestral sequence. We explored two different approaches for reconstructing the ancestral sequence: firstly, utilizing the unrooted tree returned by RAxML-NG, and secondly, rooting the tree using midpoint rooting. This second approach positions the root at the midpoint between the two longest branches. Subsequently, we compared the inferred ancestral sequences to the known ancestral sequences and calculated the Hamming distance. As a point of comparison, we repeated this process for the correct families. As expected, our findings align with previous results: the distribution of sequence similarity for SCOPer-H closely mirrors that of the correct families. Following closely are SCOPer-S and mPTP. This underscores the significant impact of method selection on downstream analysis of repertoire sequence data. Our analysis indicates that utilizing the midpoint root yields superior results for ancestral sequence reconstruction across all methods (Figure S12).

## Conclusion

With the widespread adoption of BCR repertoire sequencing, understanding the evolutionary relationships of B-cells has become increasingly feasible. However, accurate delimitation of B-cell clones is essential for any meaningful analysis. Numerous tools have been developed to tackle this challenge, employing diverse approaches. Our comparative analysis of four state-of-the-art tools revealed SCOPer-H as the optimal choice for delimiting B-cell clones in organisms with reliable reference genomes, such as humans and mice.

SCOPer-H effectively accounts for both VDJ-recombination and somatic hypermutation (SHM) utilizing a reference genome, making it well-suited for model organisms. Additionally, we found mPTP, a native phylogenetic method for species delimitation, to be effective in delimiting B-cell clones across various scenarios. Notably, mPTP does not rely on a reference genome, making it particularly valuable for analyzing non-model organisms lacking a robust reference genome. In our investigation, we concentrated on the accuracy of the tools rather than other aspects like computational time. All methods demonstrated similar processing speeds in our simulations. However, computational time might be an important consideration for analyses involving real data. Our investigation into the downstream effects of clonal assignment on ancestral sequence reconstruction revealed that the choice of clonal assignment tool significantly influences the accuracy of ancestral sequence inference. This underscores the importance of selecting the most appropriate tool for clonal family assignment, especially in the context of vaccine design and other downstream applications. In conclusion, our study provides valuable insights into the performance of different clonal assignment methods and their implications for downstream analysis in B-cell repertoire sequencing studies. These findings offer guidance for researchers in choosing the most suitable tools for their specific research goals and target organisms.

## Materials and Methods

### Simulations

To conduct a thorough analysis of the diverse tools under distinct conditions, we systematically simulated B-cell repertoires, manipulating parameters such as clone count, SHM and lineage count per clone. These simulations were executed with partis [26], a Hidden Markov Model-based framework specifically designed for B- and T-cell receptor sequence annotation. The utilization of partis in these simulations ensures a reliable and standardized platform for assessing the performance of the tools across a spectrum of conditions within the B-cell repertoire. In our study, we used the simulate-from-scratch option within partis to generate a comprehensive dataset comprising 1200 simulated B-cell repertoires. These repertoires were systematically simulated across 24 distinct parameter configurations. Specifically, we simulated 6 SHM rates: 0.001, 0.005, 0.01, 0.05, 0.1, 0.2 (mutation rate per position), encompassing a broad spectrum of mutation scenarios. The true SHM rate is estimated to be 1 in 10^3^ base pairs per cell division [2]. This is challenging to rescale for a simulation setup, primarily because the number of cell divisions per sample in nature is variable and not always known. However, by simulating a spectrum of SHM rates, we aim to capture trends in performance across all methods. Antibody sequences may typically exhibit divergence of up to 5% from their original germline sequence [2]. Hence, we calculated the extent of divergence between our simulated sequences and their true ancestral counterparts to assess the variability. Our examination indicated that SHM rates of 0.001, 0.005, 0.01, and 0.05, on average, adhere to this criterion (Figure S8). Thus, the majority of our simulations reflect a realistic degree of somatic hypermutations. This approach allows us to explore how different SHM rates impact the performance of each method and to identify the most suitable method for various scenarios. Similarly, we chose four values for the mean number of leaves per clonal family drawn from a geometric distribution. For our main simulation setup we chose to simulate 16 different clonal families. We later also explored this parameter more by choosing 10, 20 and 50 as different clone sizes. This extensive range of parameter setups ensures a robust evaluation of the tools under diverse conditions, facilitating a nuanced understanding of their performance across the specified parameters. After observing that all methods tend to oversplit families and result in many singletons, simulated clonal families consisting of one sequence, we decided to remove singletons for our analysis. Since the goal of our study is to test the method’s ability to discern families and not singletons this simplifies the analysis.

In order to further pinpoint the differences between the tested tools we conducted additional simulations without partis, but using an algorithm developed in our lab, across the same parameters. This approach enabled us to simplify the simulations, focusing specifically on scenarios where SHM exclusively impacts the junction regions, excluding the V, D, and J genes within a clonal family. This refinement was particularly motivated by the analytical emphasis of SCOPer on the junction regions of sequences. To simulate realistic sequences, we utilized the IMGT [17] reference directory, which contains most of the known human V,D, and J genes. For each naive sequence we randomly sampled from the reference genes and joined them together, adding a varying amount of N and P nucleotides in between. For the generation of SHM, targeting the junction regions within a clonal family only, we employed the phangorn package [27] in R, leveraging its simSeq function. This function facilitates the simulation of sequences based on a specified phylogenetic tree. This targeted simulation approach allowed us to craft scenarios precisely aligned with the questions and considerations specific to SCOPer’s analytical focus on junction regions.

We additionally created a separate simulation set to test how the methods perform when the sequences do not align well with the reference genomes. To achieve this, we took the original V gene sequences from IMGT and introduced three deletions and three insertions of sizes varying from 1 to 4. We also included point mutations as a parameter, ranging from 20 to 40. Due to the relatively short length of the D and J genes, we did not modify them. The rest of the simulation procedure remained the same as in the previous setup, with 6 N and P nucleotides added at each junction.

### Tools

In this study we evaluated the performance of multiple state-of-the-art tools for clonal assignment in B-cells. In the following we specify the exact parameters used for each tool in this study.

#### MiXCR

MiXCR is a tool for immune data analysis particularly favored by major pharmaceutical companies [28] for diverse downstream analyses, one of them being clone identification. It discerns the clonal families by sequence identity on specific gene features. In our study, we specifically opted to assemble clonotypes based on the CDR3 region, aligning with common practices in real-world analyses.

#### Change-O

Change-O is a toolkit with diverse applications in immunogenetics. It depends on an alignment to a reference genome for clonal family assignment. In our study, we chose to align the sequences using IMGT/V-QUEST [29], as our analysis revealed no discernible difference in results between IMGT/V-QUEST and IMGT/HighV-QUEST [12] for our purposes. To streamline this process, we adapted the vquest API provided by the ShawHahnLab [30] for the required output type excel.

#### SCOPer

SCOPer leverages the output of Change-O but diverges in its approach by incorporating SHM. Instead of discerning clonal relationships solely based on common V-gene, J-gene, and identical junction regions, SCOPer determines clonal relatedness based on a minimum similarity threshold for the junction regions. In SCOPer-H, users must define a threshold, while SCOPer-S, the spectral model, autonomously determines optimal threshold values for each subgroup identified by Change-O. For our evaluation, we adhered to the default cutoff of 0.15 for SCOPer-H, as suggested by Nouri et al. [10]. This threshold, commonly utilized in previous studies on human B-cell repertoires [25, 31, 32, 33], permits a deviation of up to two mutations in the CDR3 region among sequences within the same clonal family.

#### mPTP

mPTP is a single-locus species delimitation method which uses maximum-likelihood and Markov chain Monte Carlo sampling [15]. It takes a binary phylogenetic tree T as input. We employed RAxML-NG [16] in our study to infer the phylogenetic tree from the sequence data, aligning with the recommended methodology for implementing the mPTP approach by the authors. The objective of mPTP is to find a binary subtree G of T such that the likelihood of the branch lengths of G fitting an exponential distribution and the branch lengths of each maximal subtree of T formed by the remaining branches fitting an exponential distribution is maximized. Here, G represents the speciation process, while all other maximal subtrees of T represent the coalescent processes. mPTP uses a dynamic programming approach that traverses all nodes of T in postorder traversal. The delimitation with the smallest Akaike Information Criterion score is selected as the final result. To evaluate the confidence of the chosen delimitation, mPTP utilizes an MCMC approach.

### Metrics for assessing performance

To comprehensively evaluate the performance of all tools, we used various measures. In all our analyses, we opted to exclude singletons, which are derived clonal families containing only a single sequence, in order to reduce noise and because they are disregarded in real-life analyses as well. For a broad overview and indirect assessment, we computed the MSE of the Median Family Size. This metric serves to determine whether the identified families align closely in size with the actual families, providing valuable insights into the overall accuracy of family size assignments across the evaluated methods. To find the cause of large MSEs we also counted the number of families that the methods derived for each simulation and compared it to the real number of clonal families. This helped us to understand whether the methods were over or under splitting the clonal families.

For more direct and more interpretable metrics, we computed precision, recall, and the F1-score. In our evaluation, analogous to other studies with similar assessments [11, 34, 35, 36], we defined True Positives (TP), True Negatives (TN), False Positives (FP), and False Negatives (FN) according to the following criteria:

For each sequence *x_i_*:

TP : # of sequences from the same family as *x_i_* that are correctly identified as being in the same family TN: # of sequences from a different family as *x_i_* that are correctly identified as being in a different family FP: # of sequences from a different family as *x_i_* that are incorrectly identified as being in the same family FN: # of sequences from the same family as *x_i_* that are incorrectly identified as being in a different family

We then calculated the precision, recall and F1-score (harmonic mean of precision and recall) for each *x_i_*. Recall answers the question: Of all sequences that were clustered together, how many actually belong to the same family? Precision answers the question: Of all the sequences belonging to the same family, how many were correctly clustered together? The F1-score is the harmonic mean of precision and recall and therefore increases when reducing the instances where a single clonal family is divided into multiple groups and the cases where multiple clonal families are combined into a single group. For all 7 quantities we averaged them over all sequences to end up with a value per simulation.

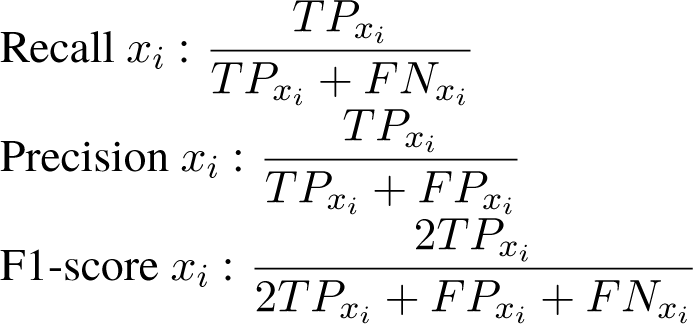

#### Ancestral Sequence

Our downstream analysis consists of the evaluation of the ancestral sequence reconstruction. For each inferred clonal family comprising more than two sequences we first aligned the sequences and subsequently constructed a phylogenetic tree using RAxML-NG. Ancestral sequences were then reconstructed using RAxML-NG with the GTR model. As input we used both an unrooted tree as returned by RAxML-NG and a tree rooted using the midpoint root, which roots the tree halfway between the longest two tips. The Hamming distance between the inferred sequence and the correct naive sequence from the simulations was calculated. As a control, we repeated this process for correct families, recognizing that a correct family does not necessarily lead to the correct ancestral sequence, owing to uncertainty in the ancestral reconstruction itself.

## Acknowledgements

We acknowledge support from NIH grant R35GM151348 to MP. We thank members of the Pennell, Edge, and Mooney labs for their thoughtful comments on this study.

## Data Availability

All the code used for the simulations and all analyses can be found at https://github.com/katalinandreavoss/ clonal-misclassification.

## Supplementary Material

**Figure S1:**
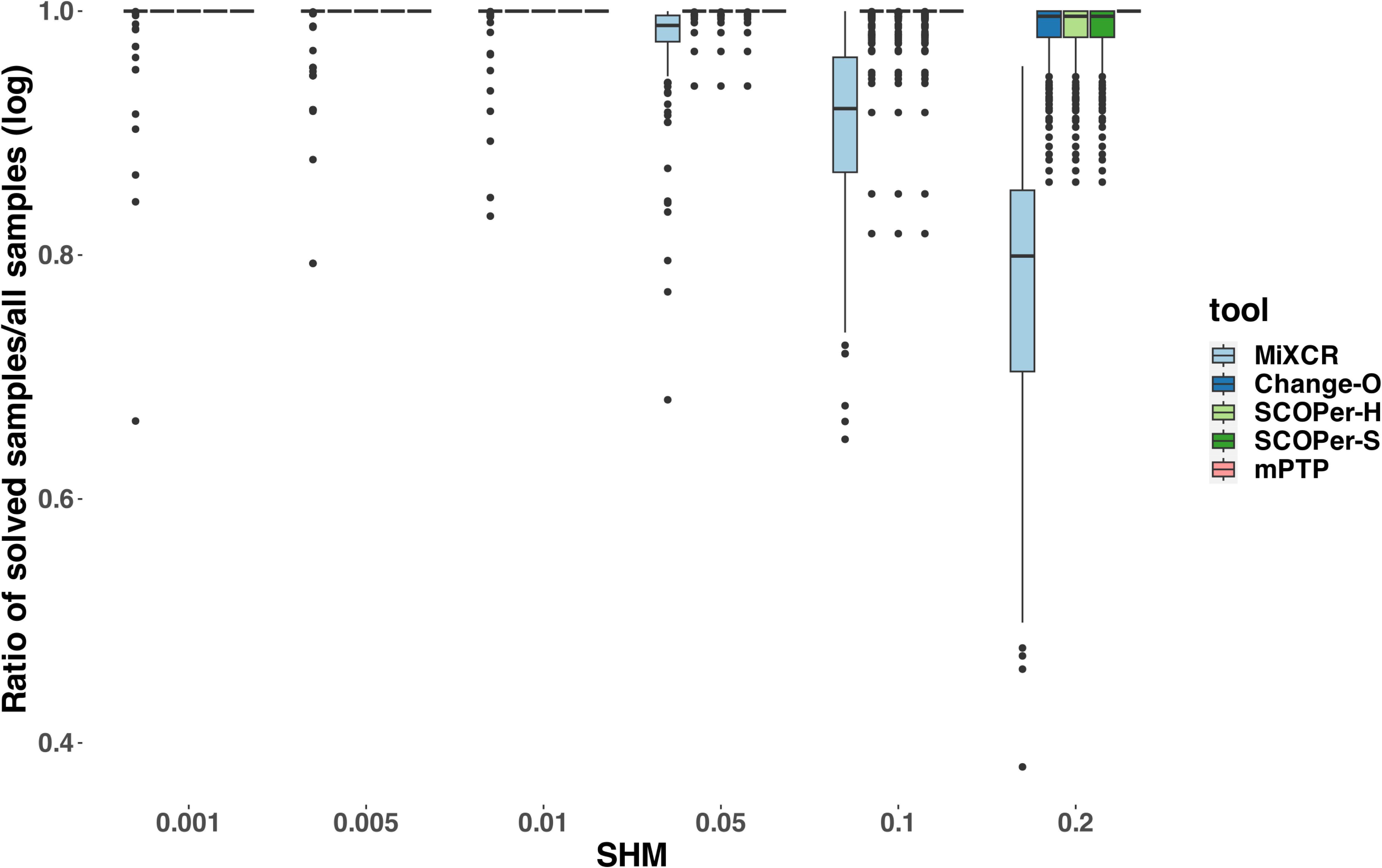
Ratio of solved samples/all samples (log) across different SHM rates. For this analysis we counted the amount of sequences that the methods assigned to a clonal family and divided that by the number of all sequences from the input.

**Figure S2:**
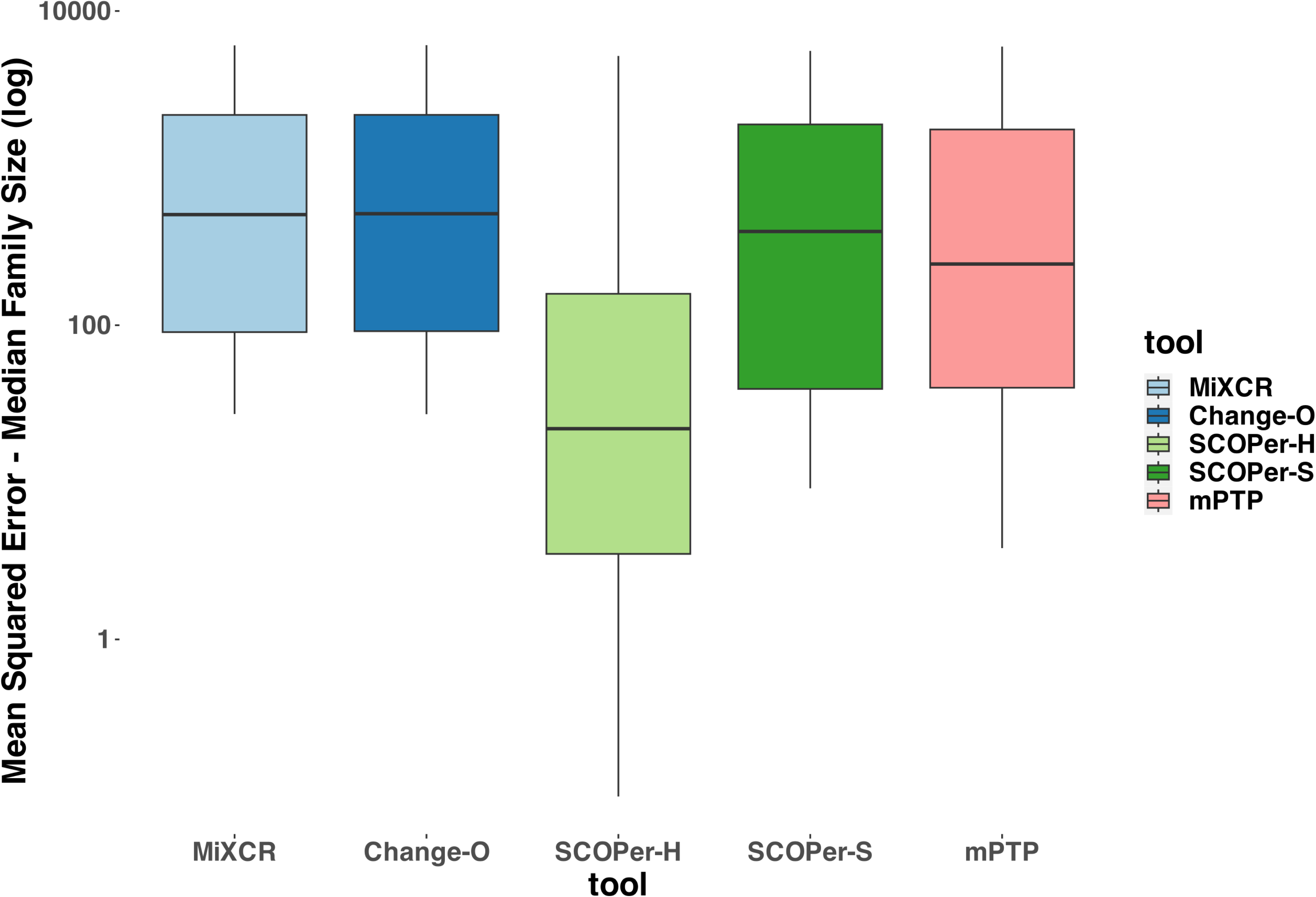
MSE of the median family size. For this analysis we removed singletons. We calculated the median family size of the true clonal families and compared them to the median family size of the derived clonal families for each method.

**Figure S3:**
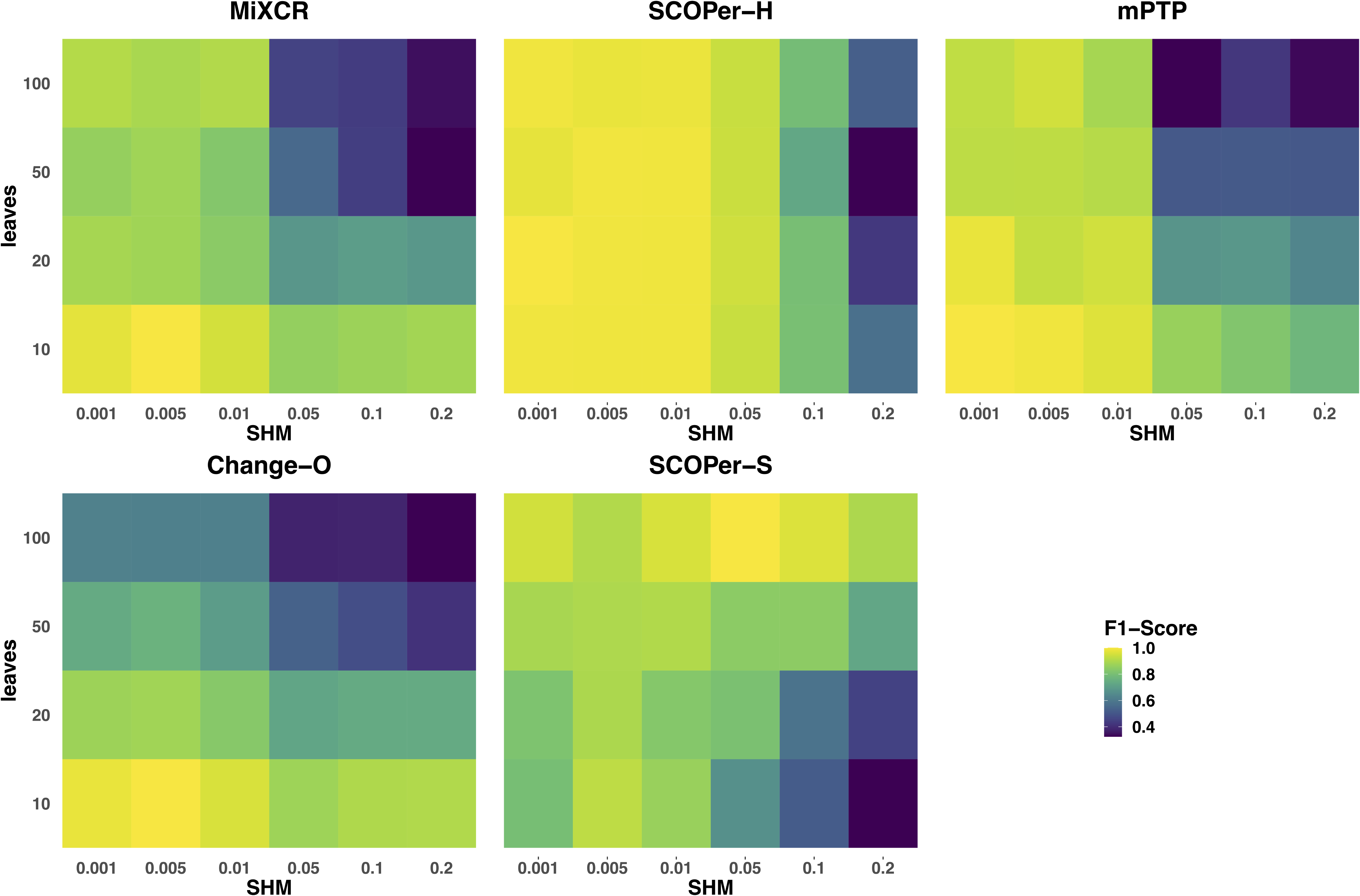
F1-Score across different parameters for all methods. For this analysis we removed singletons. The F1-score was calculated by taking the average of all simulations with the specific leaf and SHM configuration.

**Figure S4:**
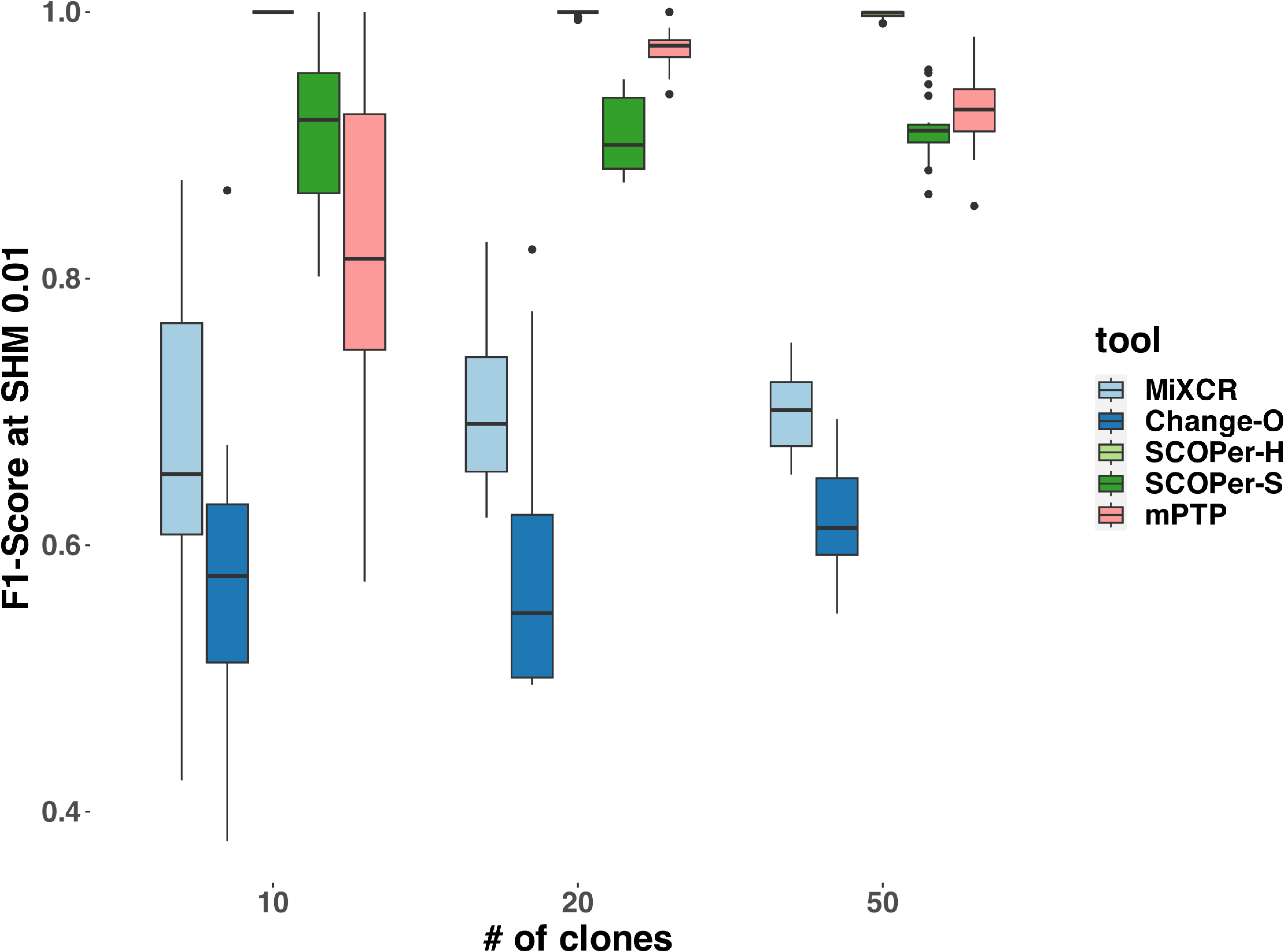
F1-Score across different numbers of clones for all methods. For this analysis we removed singletons. The F1-score was calculated by taking the average of all simulations with the same number of clones.

**Figure S5:**
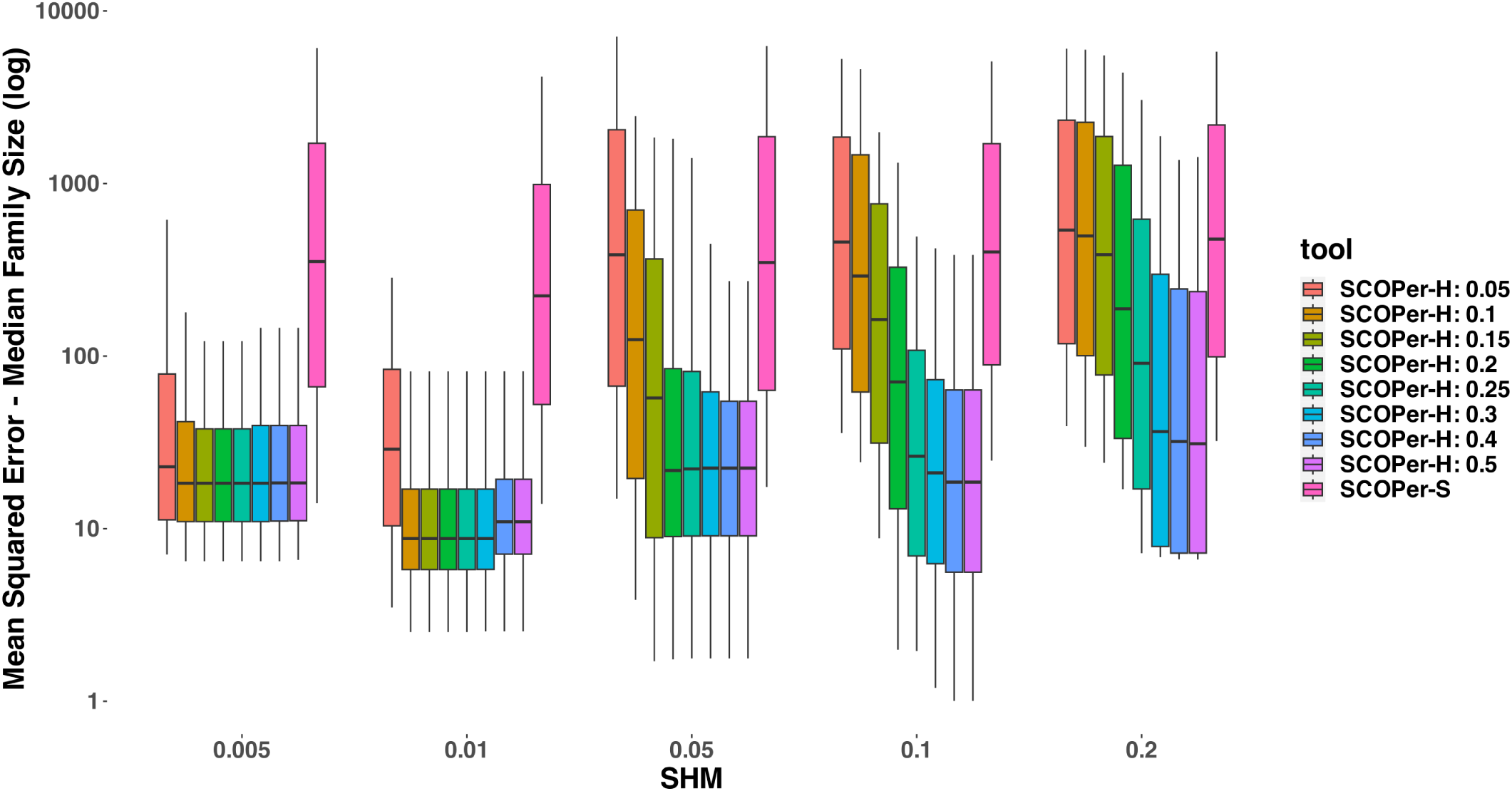
MSE of the median family size for different SCOPer-H thresholds. For this analysis we removed singletons. SCOPer-H: 0.15 is the one used in all other analyses.

**Figure S6:**
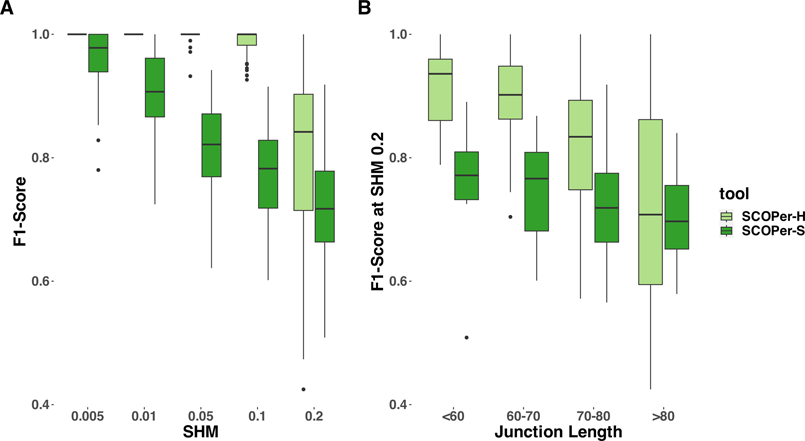
F1-Score across different parameters for SCOPer-H and SCOPer-S. For this analysis we removed singletons. A) different SHM rates B) different junction region lengths (nt). The junction region length was calculated by SCOPer.

**Figure S7:**
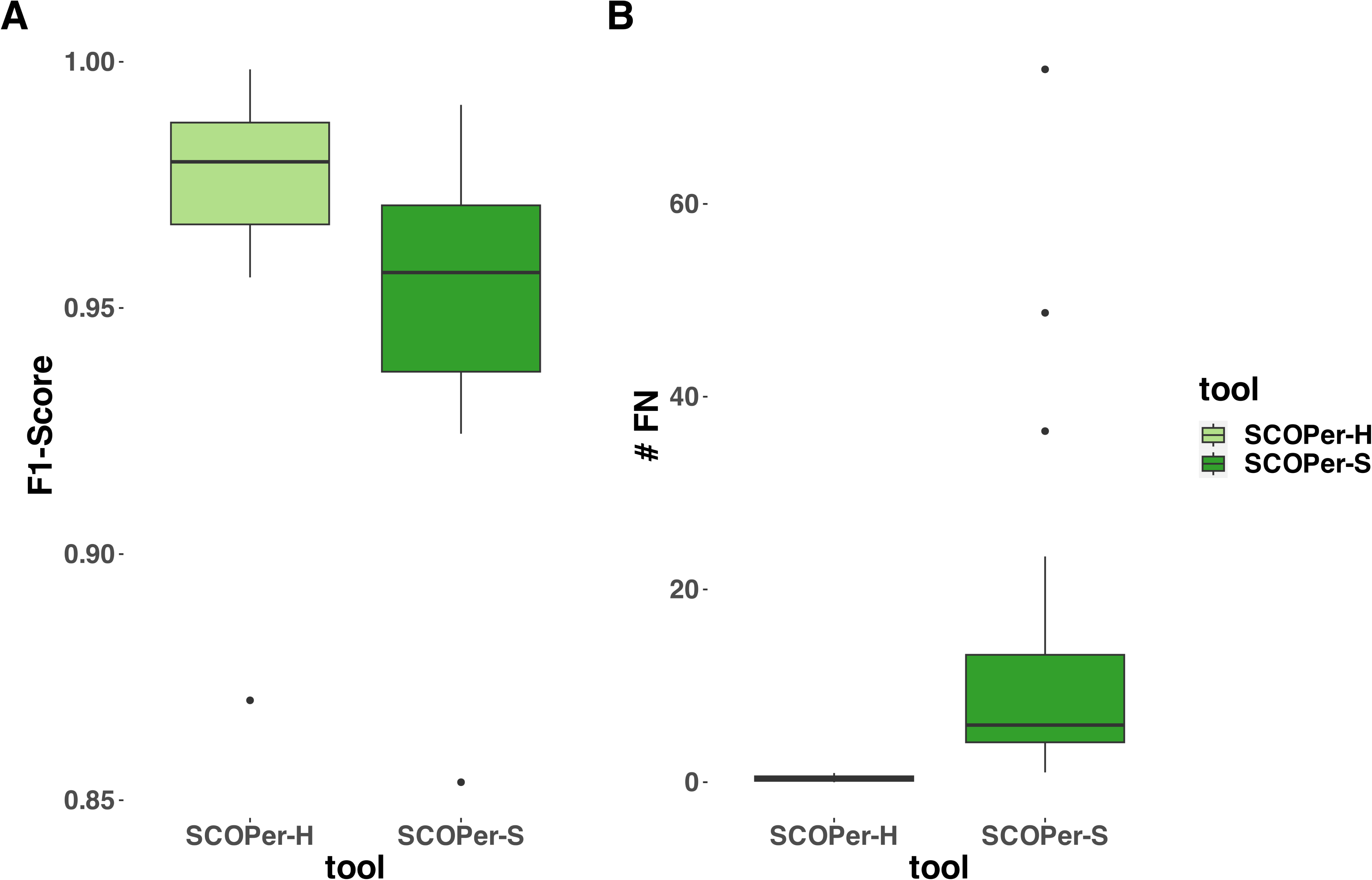
Evaluation of a subset of simulations provided by Nouri et al. [11] A) F1-score for SCOPer’s hierarchical and spectral model B) number of False Negatives for SCOPer’s hierarchical and spectral model. For each simulation we took the 20,000 first results and evaluated them.

**Figure S8:**
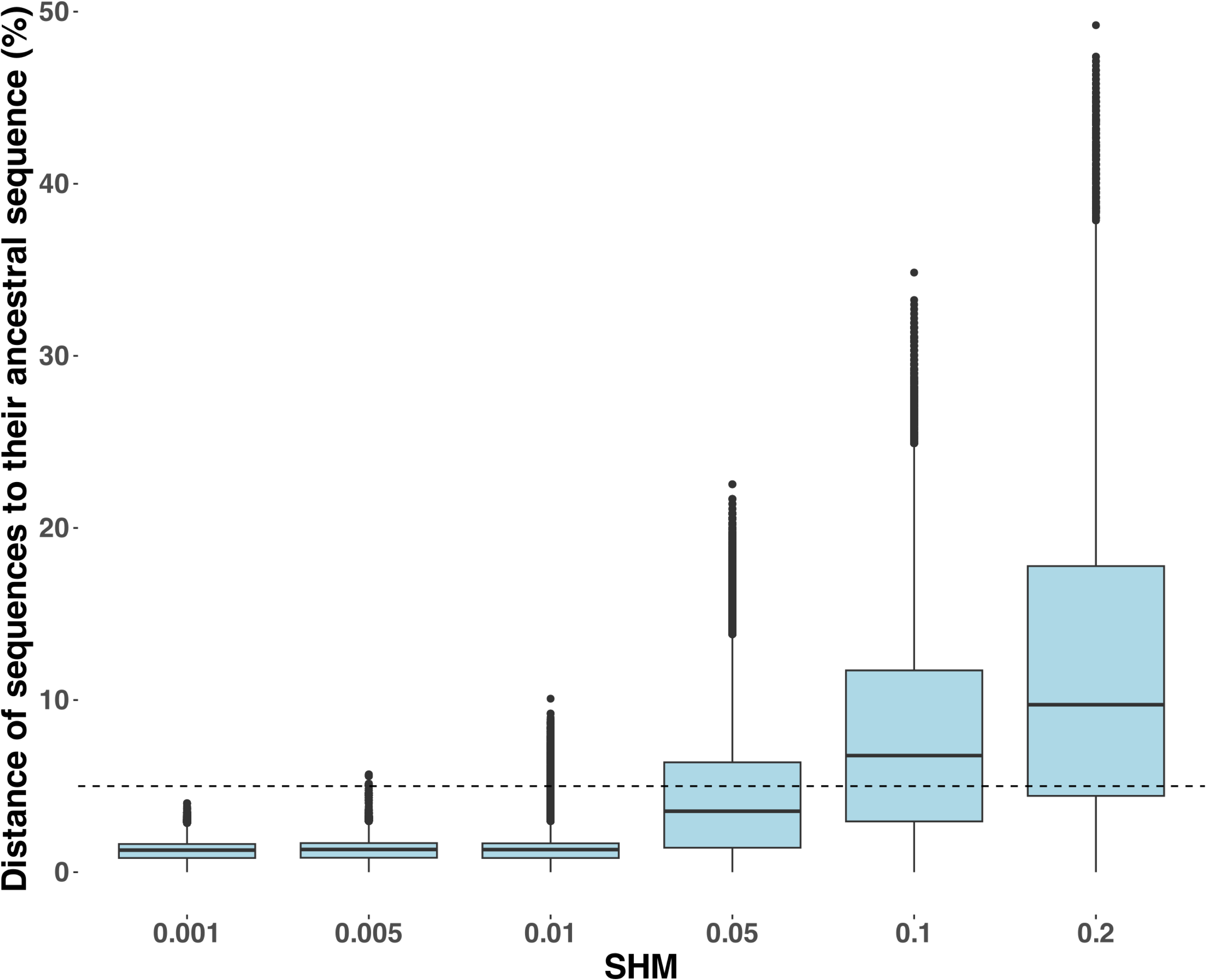
Sequence Divergence between the simulated sequences and their ancestral sequence. The dashed line is at 5%, which is a known level of divergence in real sequences.

**Figure S9:**
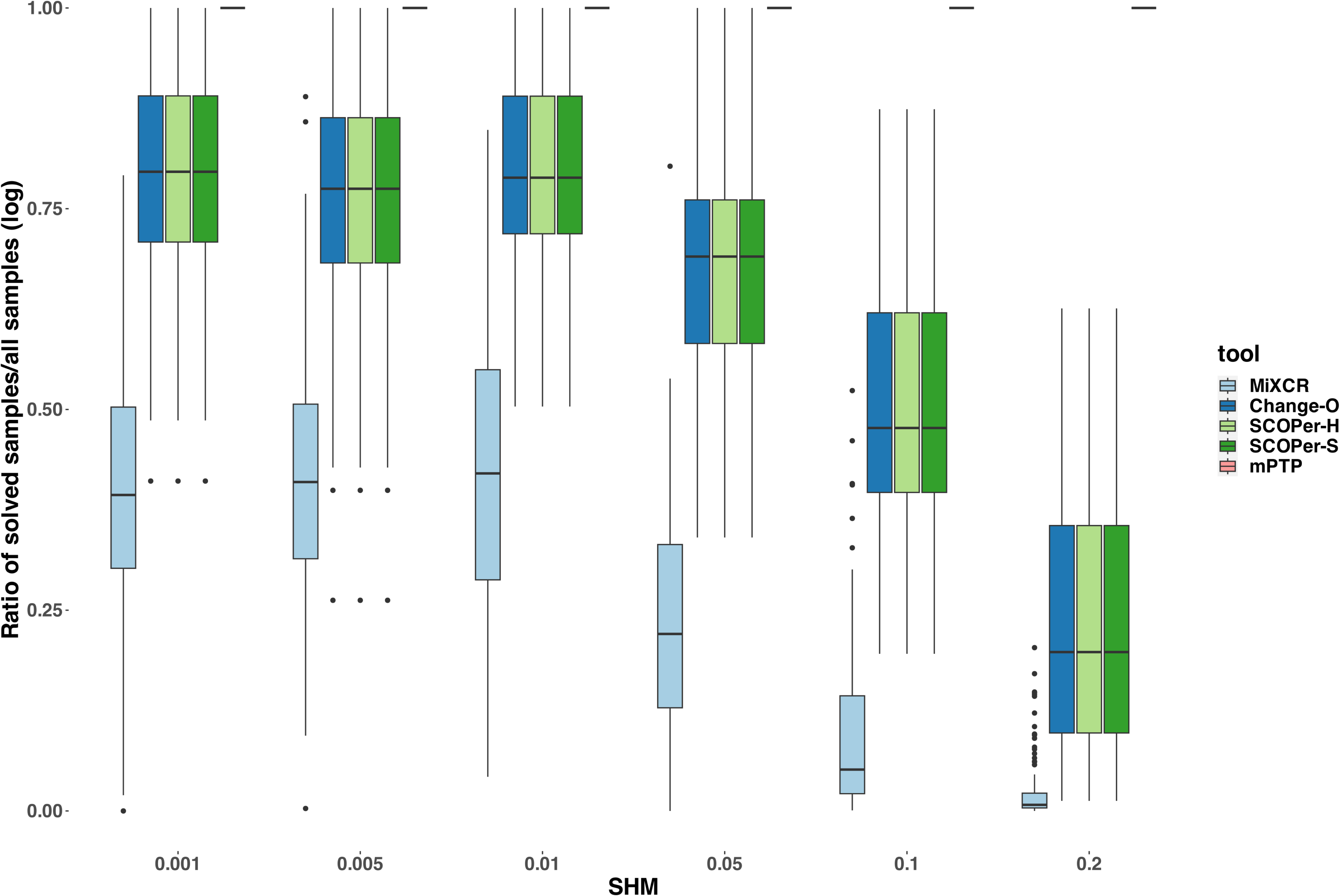
Ratio of solved samples/all samples (log) across different SHM rates in simulations with “fake” V genes. For this analysis we counted the amount of sequences that the methods assigned to a clonal family and divided that by the number of all sequences from the input.

**Figure S10:**
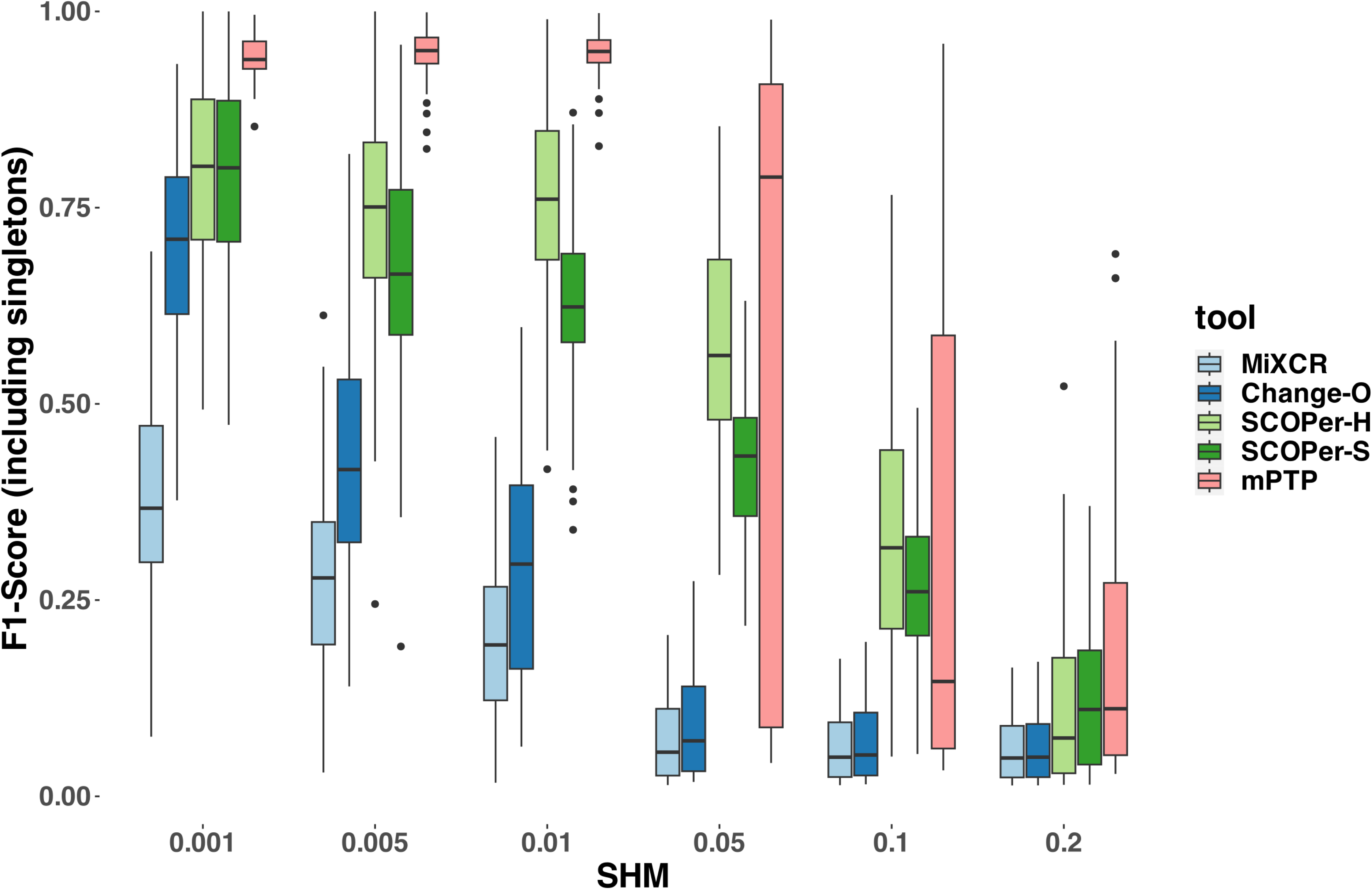
F1-Score yielded by the different methods across different SHM rates for simulations with “fake” V genes (includes singletons) The F1-score was calculated by taking the average of all simulations with the specific SHM configurations.

**Figure S11:**
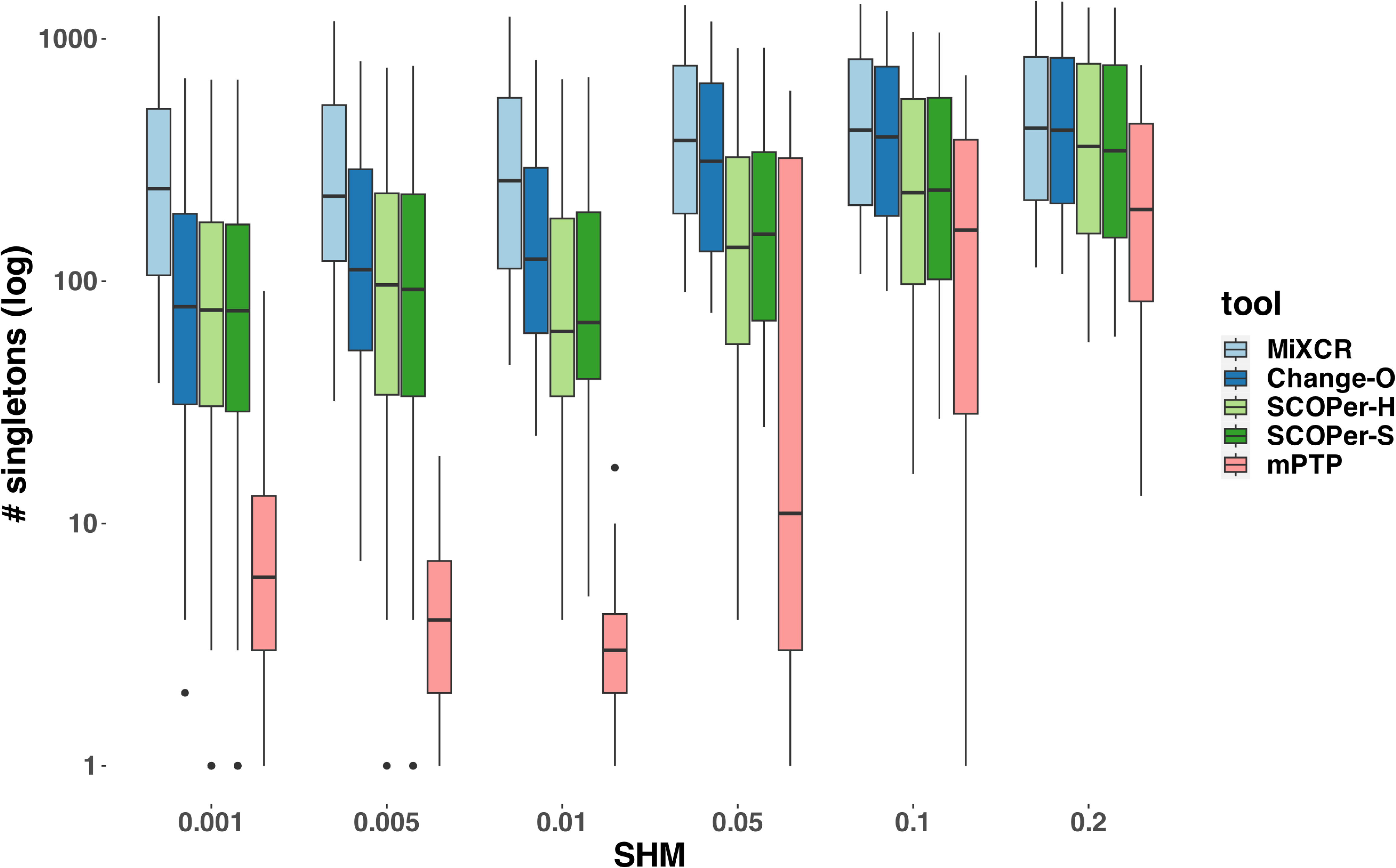
Number of singletons derived by each method for simulations with “fake” V genes

**Figure S12:**
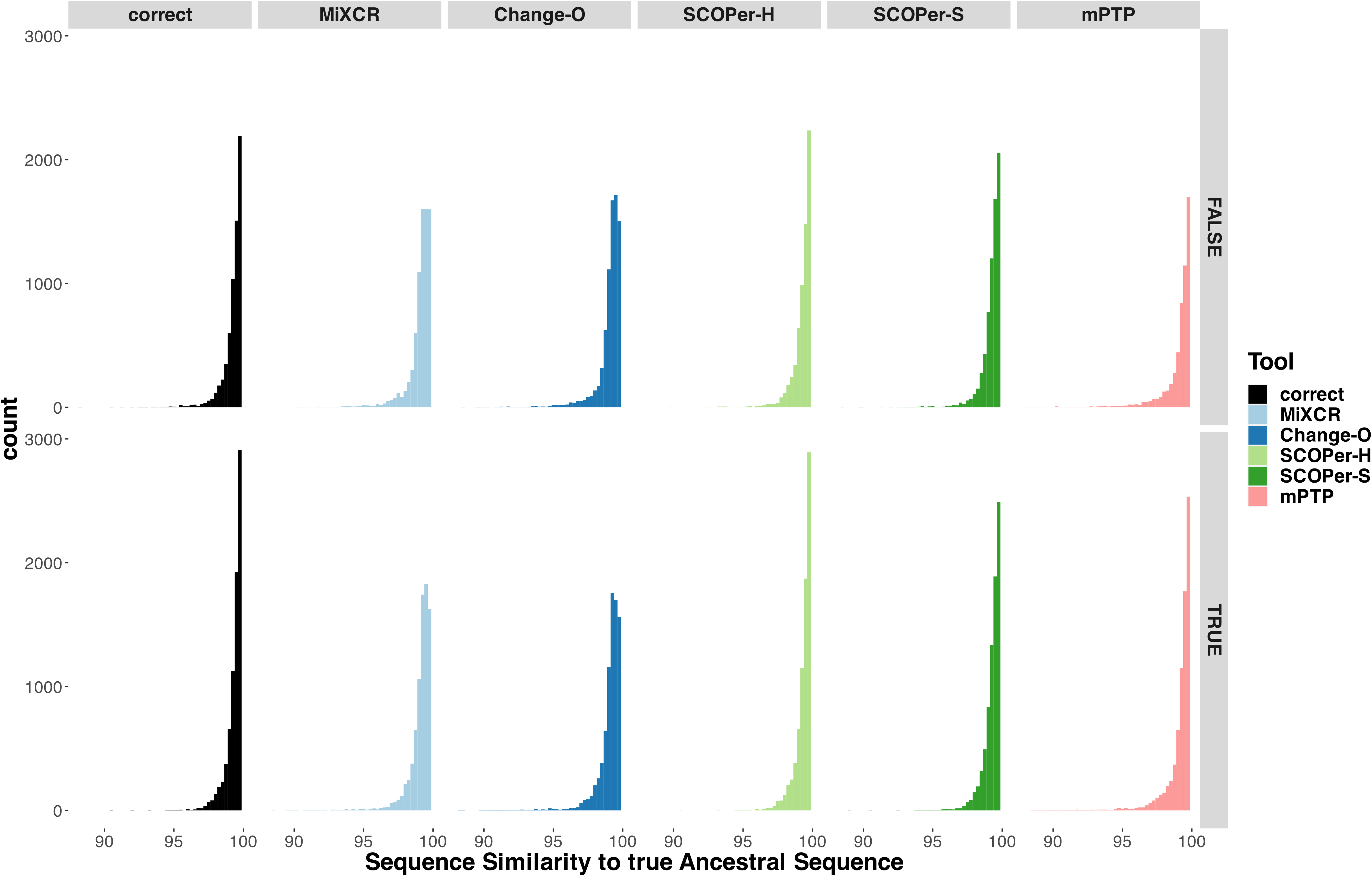
Sequence Similarity between the real ancestral sequence and the derived ancestral sequence. based on the clonal families discerned by the methods split between using the unrooted tree provided by RAxML-NG (FALSE), and rooting at the midpoint (TRUE)

